# Fungal community dynamics and carbon mineralization in coarse woody debris across decay stage, tree species, and stand development stage in northern boreal forests

**DOI:** 10.1101/2022.01.31.478531

**Authors:** Saskia C. Hart, Teresita M. Porter, Nathan Basiliko, Lisa Venier, Mehrdad Hajibabaei, Dave Morris

## Abstract

Fungi are primary agents of coarse woody debris (CWD) decay in forests, playing an essential role in nutrient cycling and carbon storage. Characterizing fungal communities within CWD will promote further understanding of the fungal controls on CWD decomposition. We compared fungal community assemblages using alpha and beta diversity metrics, carbon mineralization, and physical and chemical properties of CWD across 3 tree species (trembling aspen [*Populus tremuloides*], black spruce [*Picea mariana*]*, and* jack pine [*Pinus banksiana*]), 5 decay classes, and 2 stand development stages, differing in time-since-stand replacing disturbance (i.e., young/self-thinning and mature/steady-state) in Ontario’s boreal forest region. In total, we sampled 180 individual CWD logs from 6 independent stands, with 3 replicates per each species × decay class combination at each site. We found that fungal community structure significantly differed across tree species, decay stage, and stand age. Higher proportions of white rot fungi were found in trembling aspen CWD, whereas higher proportions of brown rot fungi were found in black spruce and jack pine CWD. Proportions of specialized wood decay fungi increased with decay stage and were higher in CWD located in mature forest stands. Fungal diversity was highest in decay class 4 CWD. We found that Mn and K concentrations, total carbon, C/N ratio, carbon mineralization (mg CO_2_ g dry CWD^-1^ d^-1^), and moisture content were important predictors of fungal composition across CWD species and/or decay stage, though how CWD chemistry influences fungal species composition (and vice versa) is unknown. Carbon mineralization was highest in trembling aspen CWD and increased with decay stage, perhaps facilitated by increased N concentrations. This study suggests that forest management guidelines that consider both deadwood quantity and quality will support a broader range of fungal species and communities through post-disturbance stand development, thereby conserving biodiversity over the longer-term in our managed forest systems.

## 1. Introduction

Saproxylic fungi are primary agents of coarse woody debris (CWD) decomposition in boreal forests (Boddy & Watkinson, 1995; Harmon et al., 2004; Russell et al., 2013). The decomposition of CWD in all of its forms (i.e., across species, size, decay state, location) by saproxylic fungi is an ecologically significant function, creating habitats for fungi, insects, birds, and small mammals, promoting forest productivity and regeneration via nutrient cycling, and playing a role in forest carbon cycles (Stevens, 1997). Nutrient and carbon turnover in CWD are determined by decomposition processes and rates, which, in turn, are influenced by air temperature, moisture availability, substrate quality, and the community structure of decomposer organisms, chiefly fungi (Boddy, 2001; Edmonds et al., 1986; Harmon et al., 2004). The interactions between fungi, CWD species, and CWD decay stage also affects the rate of carbon mineralization. For example, studies have shown higher CO_2_ production in hardwood CWD, with increasing decay stage (Gough et al., 2007; Harmon et al., 2004; Herrmann & Bauhus, 2013; Kahl et al., 2015; Pastorelli et al., 2017; Wang et al., 2002). Estimating accurate carbon flux from CWD decomposition is essential in understanding the carbon balance of boreal forests in Canada as CWD can store up to 30% of aboveground carbon (Laiho & Prescott, 1999).

There have been several studies detailing fungal community composition within CWD across northern Europe, but fungal CWD community composition in northern Canada has not been fully described (Heilmann-Clausen & Christensen, 2003; Kebli et al., 2012; Kubartova et al., 2012; Ottosson et al., 2014; Ovaskainen et al., 2010). DNA sequence-based studies of community structure have been revolutionary in documenting fungal communities within CWD in northern European forests and have reduced the visual bias of fruiting body surveys (Hoppe et al., 2016; Lindner et al., 2011; Ottosson et al., 2014; Ovaskainen et al., 2010; Ovaskainen et al., 2013, Rajala et al., 2012).

In Ontario, boreal forest stands that contain black spruce (*Picea mariana* (Mill.) B.S.P), jack pine (*Pinus banksiana* Lamb.), and trembling aspen (*Populus tremuloides* Michx.) are of economic importance as they are sources of high-quality wood and year-round harvesting opportunities (OMNR, 2003). Stand structure, including the presence and input of CWD as slash, in managed silvicultural systems is a critical component of biodiversity and so understanding the nature of CWD inputs is essential in understanding subsequent decomposition processes (OMNR, 2003). Examining the relationship between saproxylic fungi and CWD will assist in evaluating the effectiveness of current stand-level management approaches applied to maintain sufficient CWD in forest stands (Berch et al., 2011). Our objective is to compare fungal community structure and function using high-throughput sequencing with respect to CWD tree species, decay class, and stand development stage.

Our analysis focused on two key questions. The first question was: *how does the fungal community composition in CWD change across tree species, decay class, and stand development stage?* Here we predicted that fungal community composition would differ (i.e., increased proportions of brown rot versus white rot fungi) between conifer (black spruce or jack pine) and hardwood (trembling aspen) due to differences in wood chemical and physical properties. Fungal species richness and community composition in CWD has been shown to be strongly driven by tree species due to the differences in physical and chemical properties of wood (Baber et al., 2016; Boddy, 2001; Hoppe et al., 2016). Fungi differ by CWD species primarily in their ability to degrade lignin. White rot fungi have the ability to break down lignin via ligninolytic enzymes and are largely associated with hardwood CWD, whereas brown rot fungi cannot break down lignin and are largely associated with conifer CWD (Schwarze, 2007). We also predict greater fungal diversity as wood decay advances due to increased niche availability (Heilmann-Clausen & Christensen, 2003). We predict differences in fungal community structure and diversity (i.e., greater diversity in the self-thinning stands compared to the mature/overmature stands), as a function of stand age (stand development stage) due to top-down controls of canopy density and likely differences in throughfall (i.e., volume of intercepted rainfall, and chemical modification as rainwater moves down through the canopy), and litterfall inputs to the forest floor (Hawkes et al., 2011; Hu et al., 2021; Laperriere et al., 2019; Ryan et al., 1997). Recent literature has supported the pattern of increased species diversity and richness with increasing decay class, suggesting that niche availability may influence fungal diversity and richness, along with other external factors, such as nutrient dynamics of CWD (i.e., import/immobilization versus release), CWD species, and climate (Arnstadt et al., 2016; Baldrian et al., 2016; Kurbartová et al., 2012; Rajala et al., 2012; Yamashita et al., 2015).

Our second question was, *what are the relationships between fungal communities and the physical and chemical properties of CWD, and C mineralization (i.e., rates of microbial CO_2_ production) in CWD?* Here we predicted that there will be significant differences between fungal community composition and activity (within-log CO_2_ production) for those communities that have become established in hardwood versus conifer tree species, with increases in specialized wood decay fungi in each species as decay advances. We predicted these changes are linked to changes in both physical (i.e., wood density) and chemical (i.e., C/N ratios, base cation concentrations) properties. In a European study comparing fungal community structure in CWD between European beech (*Fagus slyvatica* L.) and Norway spruce (*Picea abies* (L.) Karst.), decay class, relative wood moisture, pH, wood volume, wood density, C/N ratio, and lignin concentration all correlated significantly to the variation in fungal community structure (Hoppe et al., 2016).

## 2. Materials and methods

### 2.1 Site Descriptions

A total of 6 fire-origin sites were included in the study. Three sites were located 30km northeast of Thunder Bay, Ontario, with the other three sites located 30km northeast of Chapleau, Ontario (Figure 1). Although the site clusters (3 sites in northwestern Ontario, and 3 sites in northeastern Ontario) were separated geographically, the published long-term climate normals for these locations overlapped. For example, the northeastern Ontario sites have experienced average annual precipitation rates ranging from 652mm to 1029mm and an average annual temperature range from −0.5°C to 2.5°C (Crins et al., 2009). Comparatively, the northwest Ontario sites have experienced an average annual precipitation between 654mm to 879mm, and an average temperature range from −1.7°C to 2.1°C (Crins et al., 2009). All sites were generally conifer-dominated mixedwoods, varying in proportions of black spruce, jack pine, and trembling aspen, with northwest sites primarily dominated by black spruce and northeast sites primarily dominated by jack pine (Table 1). Soil textures were generally coarse to fine loams, with moderately dry to fresh moisture regimes (Table 1). In terms of stand ages, the northwest sites were mature to over-mature stands, ranging in age from 84-109 years that contrasted with the younger northeast sites (53-66 years old) that were still in the self-thinning stand development stage (Table 1).

**Figure 1:**
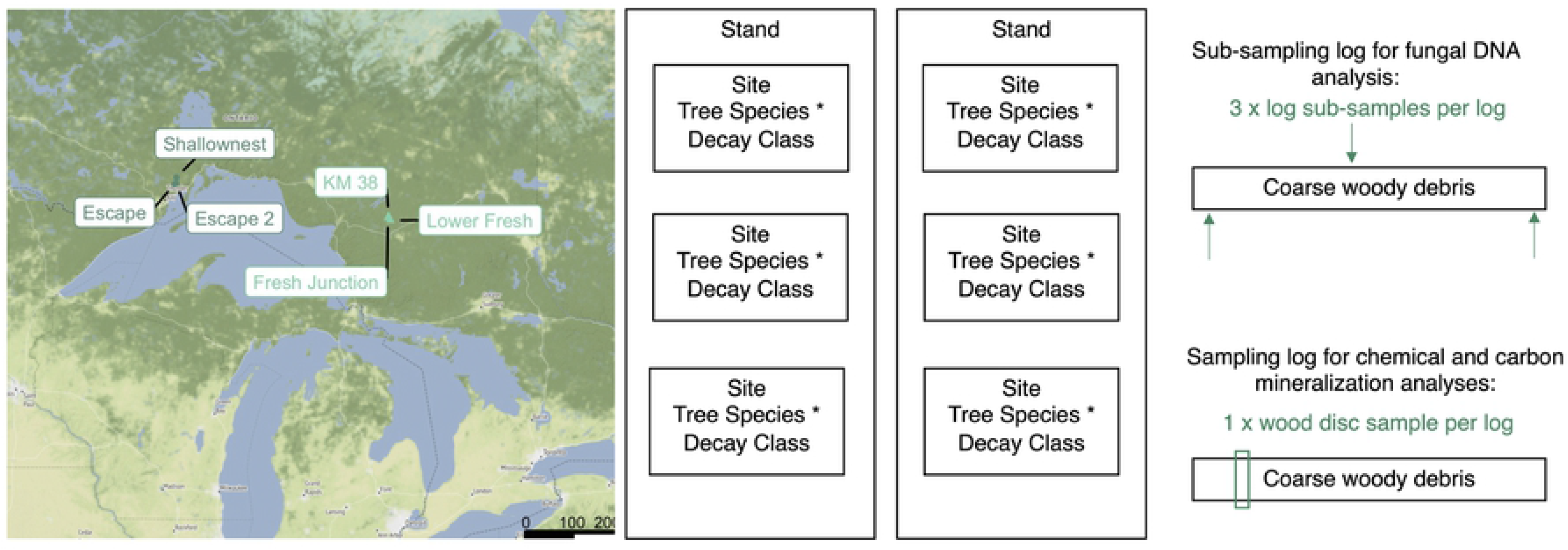
Strategy for sampling coarse woody debris (CWD) logs for analysis. Samples taken across 2 stands (mature, self-thinning), 6 sites (Shallownest, Escape, Escape 2, KM 38, Fresh Junction, and Lower Fresh), 2 wood types (softwood [black spruce or jack pine], hardwood [trembling aspen]), and 5 decay classes (1, 2, 3, 4, 5). 180 CWD logs were sampled in total (30 per site across decay class and wood type). Mature sites are labelled with circles and self-thinning sites with triangles. Scale bar shows distance in kilometers. Sites were plotted using the ‘ggmap’ package with terrain stamen maps (Map tiles by Stamen Design, under CC BY 3.0. Data by OpenStreetMap, under ODbL. Available from http://maps.stamen.com/#watercolor/12/37.7706/-122.3782) (Kahle & Wickman, 2013).

**Table 1:** Site descriptions of sampled plots in both self-thinning and mature sites in northern Ontario, Canada. Abbreviations for tree species are as follows: black spruce (Sb), jack pine (Pj), trembling aspen (Po), balsam fir (Bf), and white (paper) birch (Bf).

| Site Name | Stand Development Stage | Moisture Regime | Dominant Soil Texture | Species Composition | Age |
| --- | --- | --- | --- | --- | --- |
| Shallow Nest | Mature | 2 | Fine Sandy | Sb<sub>70</sub>Pj<sub>30</sub> | 109 |
| Escape | Mature | 4 | Sandy Loam | Sb<sub>50</sub>Po<sub>20</sub>Pj<sub>10</sub>Bf<sub>10</sub>Bw<sub>10</sub> | 99 |
| Escape 2 | Mature | 2 | Sandy Loam | Pj<sub>40</sub>Po<sub>30</sub>Bw<sub>20</sub>Sb<sub>10</sub> | 84 |
| KM 38 | Self-thinning | 3 | Coarse Loamy | Pj<sub>50</sub>Po<sub>40</sub>Bw<sub>10</sub> | 53 |
| Junction | Self-thinning | 1 | Coarse Sandy | Pj<sub>70</sub>Po<sub>20</sub>Sb<sub>10</sub> | 61 |
| Lower Fresh | Self-thinning | 2 | Fine Loamy | Pj<sub>100</sub> | 66 |

### 2.2 Coarse Woody Debris Sampling

All sampling of *in situ* downed (in full contact with forest floor) coarse wood tree stems occurred in June 2017. Selection of tree species (hardwood versus conifer) was based on the existing dominant tree species at each site. This resulted in the sampling of trembling aspen coarse woody debris (CWD) as the hardwood species across all sites. For the conifer sampling, jack pine was the dominant species of CWD sampled on the three self-thinning stands and one of the mature sites. The last two mature, northwestern sites were dominated by black spruce. CWD log size was selected using the following criteria: a minimum diameter of 7cm on at least one end of the log, and a minimum length of 150cm. CWD pieces averaged 186cm in length and 11.2cm in diameter. For all cases, the existing CWD was classified into 5 decay classes (Table S1). At each of the 6 sites, sampling included 3 hardwood and 3 conifer replicates for each of the 5 decay classes, resulting in the sampling of 30 CWD logs at each site (Figure 1).

For fungal community analysis, wood samples were taken from each log by drilling a hole using a 3/8” drill bit and collecting the wood shavings in a plastic bag. Drill was sterilized with 70% ethanol and flamed between each sample. Samples were transported to the laboratory on ice before being stored at −20°C until processed. Wood samples were taken from 3 locations on the log: one from the top-center of the log, one from the bottom-right of the log, and one from the bottom-left of the log (Figure 1). Drill holes were evenly spaced and placed in areas most representative of the log’s decay stage. Bark was scraped off prior to drilling. A total of 537 wood samples were collected across all sites (179 logs), as 1 CWD log replicate (conifer, DC4) could not be located on one of the sites (Escape).

### 2.3 Chemical Analyses of Coarse Woody Debris

Wood discs (approximately 2-3cm in width) were cut from each CWD log (approximately 20cm in from the large diameter end) across all sites in July 2016 for wood density calculations and analysis of wood chemistry (Figure 1). A total of 180 discs were collected across all sites (Figure 1). Samples were oven-dried at 50°C until constant weight was achieved, and volume was determined using diameter and width measurements from each cookie and subsequently used to calculate density.

Each sample was separated in bark, solid wood, and decaying wood components, with the dry weight mass measured for each component. Subsamples (approximately 10g) of each component were then finely ground using a Wiley Mill through a 20-mesh sieve (Thomas Scientific, Swedeboro, NJ). Chemical analyses of the individual component samples were done for total carbon (C), total Kjeldahl nitrogen (TKN), and concentrations (mg kg^-1^) of phosphorous (P), potassium (K), calcium (Ca), magnesium (Mg), and manganese (Mn) at the Ontario Forest Research Institute (OFRI) laboratory in Sault Ste. Marie, ON. Log-level (mass weighted, by component) concentrations were subsequently calculated. P, K, Ca, Mg, and Mn analysis was performed by ICP spectroscopy following a selenium digest, a modified Kjeldahl-Gunning digestion process, with selenium oxide and sulphuric acid (Varian Liberty II, Agilent Technologies Canada, Mississauga, ON). TKN analysis was performed by automated wet chemistry using the same Selenium Digest protocol, adjusting the sulphuric acid concentration from 6.5% to 4% (Astoria Analyzer, Astoria-Pacific International, Clackamas, OR). Carbon analysis was performed using the Dumas dry combustion method via a thermal conductivity detector (Elementar vario MAX CNS, Elementar Analysensyteme GmnH, Hanau, Germany). A blank and primary quality control standard were run at the beginning of each batch to check for contamination, a secondary quality control standard was run every 20 samples and a duplicate every 10 samples to check accuracy, and both a primary and secondary controls were run at the end of each batch. For any values that were “below detection”, a midpoint between 0 and the detection limit for that element was used in subsequent calculations.

### 2.4 Microbial C mineralization (CO_2_ production) of Coarse Woody Debris

Wood discs (approximately 2-3cm in width) were cut from each CWD log (approximately 20cm in from the large diameter end) across all sites in October 2018 and frozen at −20°C until processed (Figure 1). A total of 180 discs were collected across all sites (the originally missed conifer DC4 from the Escape 1 site was accounted for in the subsequent sampling) (Figure 1). Each disc was quartered, and one quarter (approximately 30g) was retained. Each sample was defrosted at room temperature (20°C), weighed, and incubated in a 200mL glass jar with a lid fitted with a sleeve stopper at room temperature (20°C) for 7 days. Syringes were used to collect 15mL of air within jars on days 1, 3, 5, and 7, replenishing the headspace with the same volume of room air for which CO_2_ concentrations were known to maintain pressure in the jars. Samples were analyzed using an SRI 8610C-0040 Greenhouse Gas model gas chromatograph (SRI Instruments, Torrence, CA) and PeakSimple Data Systems (SRI Instruments, Torrence, CA) and CO_2_ peak areas were recorded. Calibration standards of 0.1% CO_2_ (ProSpec by Praxair, Sudbury, ON) were run every 15 samples. Samples were oven-dried at 60°C for 24h and dry weight was determined. Moisture content was calculated using wet and dry weights. Rates of CO_2_ production (C mineralization) were calculated as the increase in mass of CO_2_ over time (adjusted for losses during sampling) calculated from vol vol^-1^ concentration values using the ideal gas law and standardized per dry mass of wood discs.

### 2.5 DNA Extraction and PCR amplification

DNA was extracted from log sub-samples (n=537) using the PowerSoil DNA Isolation (Mo Bio Laboratories, Inc., San Diego, California, U.S.A) according to the manufacturer’s instructions, with the following adaptations to the protocol: a 10-minute homogenization step was performed using a Mini-Beadbeater-24 (Biospec Products Inc., Bartlesville, OK) rather than the Vortex Adaptor described. DNA was quantified using the Qubit™ dsDNA BR Assay kit (Invitrogen by ThermoFisher Scientific, Eugene, OR) with a Qubit™ 4 Flurometer (ThermoFisher Scientific). Some decay class 1 and 2 samples had DNA concentrations too low for sequencing even after trying to optimize the PCRs. As a result, a sub-set of decay class 1 (n=36) and 2 (n=36) samples were selected for re-extraction, selecting for samples with the highest DNA concentrations. The subset of decay class 1 and 2 samples, plus all decay class 3 (n=108), 4 (n=105), and 5 (n=108) samples were re-extracted using the DNeasy PowerPlant Pro (Qiagen, Hilden, Germany). DNA was extracted according to the DNeasy PowerPlant Pro manufacturer’s instructions, with the following adaptations to the protocol: a 2-minute homogenization step was performed using a Mini-Beadbeater-24 (Biospec Products Inc., Bartlesville, OK) rather than the Vortex Adaptor or PowerLyzer 24 Homogenizer described, and 250μL of Solution IR, for removal of PCR inhibitors, was used instead of 175μL, as recommended by Qiagen for problematic samples. DNA was eluted in 25μL of elution buffer and stored at −20°C. Illumina adapter=tailed (italics) fungal specific primers, forward primer fITS7 (bold) (5’ *TCGTCGGCAGCGTCAGATGTGTATAAGAGACAG***GTGARTCATCGAATCTTTG** 3’), forward primer fITS9 (bold) (5’ *TCGTCGGCAGCGTCAGATGTGTATAAGAGACAG***GAACGCAGCRAAIIGYGA** 3’), and reverse primer ITS4R (bold) (3’ *GTCTCGTGGGCTCGGAGATGTGTATAAGAGACAG***TCCTCCGCTTATTGATATGC** 5’), were used to amplify the Internal Transcribed Spacer (ITS) region of fungal DNA using polymerase chain reaction (PCR) (Ihrmark et al., 2012). These two sets were chosen for their fungal specificity, as well as their ability to overcome known primer bias, to try to capture the most diverse group of fungi from our CWD samples (Bellemain et al., 2010). Fungal DNA was amplified using 2 separate primer combinations for each sample: fITS7 and ITS4R, and fITS9 and ITS4R. The PCR cocktail was set up as a 75μL reaction mixture containing 7.5μL fungal DNA, 1.5μL each of 10μM forward and reverse primer, 37.5μL HotStarTaq *Plus* Master Mix (Qiagen, Hilden, Germany), and 27μL nuclease free water that was aliquoted into three 25μL reactions. Amplification was performed using a thermal cycler program with an initial denaturation at 95°C for 5 min, followed by 30 cycles of denaturing at 95°C for 30 sec, annealing at 50°C for 30 sec, and extension at 72°C for 1min, with a final extension at 72°C for 5 min. Positive controls (DNA obtained from known fungi) and negative controls (nuclease free water) were included in every set of PCRs performed. PCR products were analyzed by electrophoresis using 1% agarose gel with TAE buffer and GelRed (Biotium, Fremont, CA) at 120V for 45 min. Samples were visualized under UV light and measured using GeneRuler 100bp plus DNA Ladder (ThermoFisher Scientific, Ottawa, ON). Samples amplified using the fITS7 and ITS4R primer set were measured on the gel as 240bp, and samples amplified using the fITS9 and ITS4R primer set were 460bp

For each sample, the triplicate PCR products were pooled into one sample and purified with the MinElute PCR purification (Qiagen, Hilden, Germany) kit according to the manufacturer’s instructions, and eluted in 10μL of nuclease free water. The 3 log sub-samples taken from 3 locations on each CWD log were further pooled, resulting in 1 purified sample per piece of CWD (n=131). Purified samples were then quantified using the Qubit™ dsDNA BR Assay kit (Invitrogen by ThermoFisher Scientific, Eugene, OR) with a Qubit™ 4 Flurometer (ThermoFisher Scientific). Equimolar amounts of each quantified marker (fITS7 and ITS4R, and fITS9 and ITS4R) were pooled together for sequencing. Samples were indexed using adapters from two different kits, TG Nextera Index Kit v2 Sets A and B (Illumina Biotechnology Co., San Diego, CA). Samples were sequenced at the Biodiversity Institute of Ontario (Centre for Biodiversity Genomics, University of Guelph, ON, CA) on an Illumina MiSeq platform using a MiSeq Reagent Kit v3 (3 300-cycle) (Illumina Biotechnology Co., San Diego, CA) with 10% of PhiX used as a control. 129 samples were successfully sequenced.

### 2.6 Bioinformatic analysis

Illumina MiSeq reads were processed using SCVUIU v1.1, an ITS metabarcode pipeline available at https://github.com/terrimporter/SCVUIU_ITS_metabarcode_pipeline and this is briefly described below. Raw paired-end reads were merged with a minimum Phred quality score of 20 at the ends and a minimum overlap of 25bp. The fITS9 forward primer was trimmed first using CUTADAPT v2.3 (Martin, 2011), setting the minimum read length to 150bp after trimming, a minimum Phred quality score to 20, and no ambiguous bases were allowed when matching primers. Untrimmed reads were then used as input to trim the fITS7 primer using the same settings as before except that up to 3 ambiguous bases were allowed. The ITS4R reverse primer was then trimmed using same settings as above with no more than 3 ambiguous bases allowed. Reads were dereplicated using VSEARCH v2.11 (Rognes et al., 2016) and only unique sequences were retained. Reads were denoised and exact sequence variants (ESVs) were generated using USEARCH v10.0.240 with the unoise2 algorithm (Edgar, 2016), removing any putative chimeric sequences, sequences with predicted errors, and globally rare sequences (Table S2). An ESV x sample table was created with VSEARCH. The ITS2 region was isolated by removing partial ribosomal RNA gene sequences with the ITSx extractor (Bengtsson-Palme et al., 2013). ITS taxonomic information was assigned with the Ribosomal Database Classifier, using the UNITE fungal ITS reference dataset 07-04-2014 (Nilsson et al., 2019; Wang et al., 2007). Merged taxonomic assignment and VSEARCH ESV tables were generated, removing rare singletons and doubletons from each sample, and was either imported into Microsoft Excel (2011) or used directly in R 3.6.0 (R Core Team, 2019) for further data analyses. Raw paired-end Illumina reads were submitted to the National Center for Biotechnology Information (NCBI) Short Read Archive (SRA) under BioProject ID PRJNA726337.

**Table 2:**
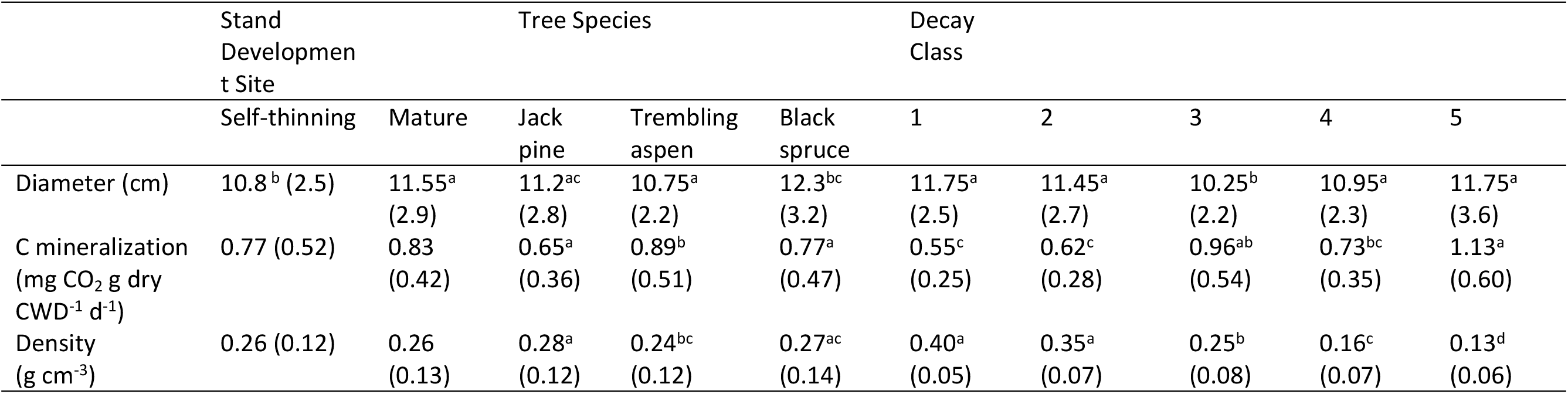
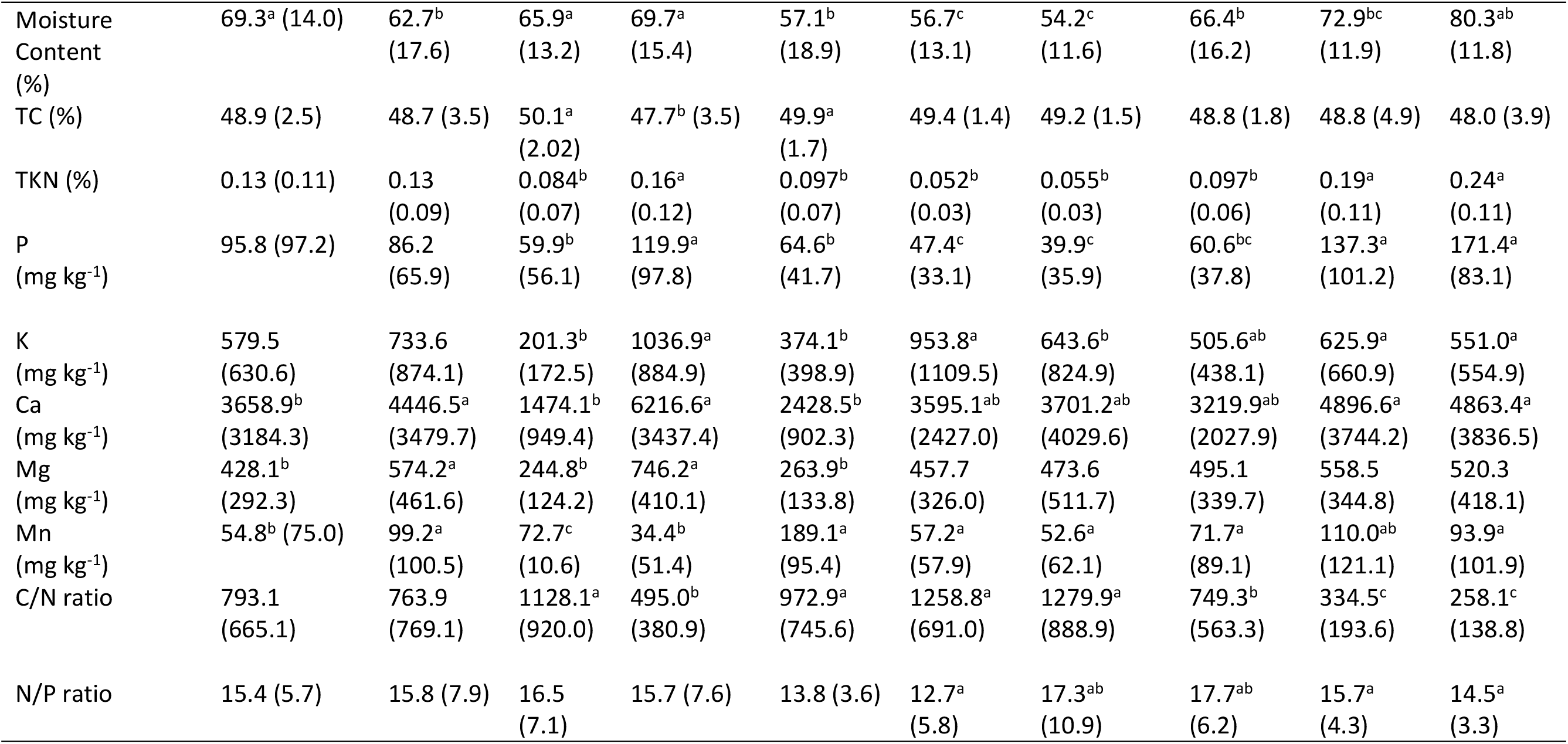
Mean physical and chemical parameters of coarse woody debris across stand development stage, tree species, and decay class (standard deviation in parentheses). Values within rows sharing the same letter do not differ significantly as determined by multiple comparisons using Tukey’s HSD test (p<0.05) (n=180).

Taxa were confidently identified at each rank across all ESVs (stand development stage, site (within stand development stage), decay class, tree species) using a 50% bootstrap support cut-off for segments <250bp (Claesson et al., 2009; Liu et al., 2012; Porras-Alfaro et al., 2014). Rarefied data was reformatted for functional trait assignment using a custom R script and processed using the open annotation tool, FUNGuild (Nguyen et al., 2016).

### 2.7 Statistical Analyses

#### 2.7.1 Wood Chemistry and Carbon Mineralization

All analyses were performed using R 3.6.0 (R Core Team 2019). The influence of stand development stage, site (within stand development stage), tree species, and decay class on wood chemical and physical properties and C mineralization was analyzed as a three-way randomized block design with replication (sites treated as blocks nested within stand development stage) using analysis of variance (ANOVA) on all CWD pieces (n=180).

Distribution of residuals for each wood physical and chemical parameter was visually assessed for normality. To determine differences within stand development stage, site (within stand development stage), decay class, and tree species, a Tukey’s HSD multiple comparison one-way analysis of variance with a threshold of p<0.05 was utilized. Chemical and physical properties were pooled across replicate CWD pieces (stand development stage, site, tree species, and decay class, and were retained for Akaike Information Criterion (AICc) modelling (n=59).

#### 2.7.2 Fungal Community Analyses

All community analyses were completed using functions from the R package “vegan” unless otherwise specified (Oksanen et al., 2019). Rarefaction curves were generated to assess read coverage per sample, and data was normalized to the 15^th^ percentile of library size using the *rarecurve* and *rrarefy* functions, respectively (Figure S1). For statistical analyses, samples (n=129) were pooled across replicate CWD pieces (stand development stage, site, tree species, and decay class) and 3 samples with less than 5^th^ percentile of reads were removed from community analyses (n=59).

Shannon-Wiener (H) diversity (Shannon & Weaver, 1949) indices were calculated for all ESVs across stand development stage, decay class, and tree species using the *diversity* function. ESV richness (S) was calculated using the *specnumber* function. Pielou’s evenness (J) (Pielou, 1966) was calculated using the equation: J=H’/log(S). The influence of stand development stage, site (within stand development stage), tree species, and decay class on ESV richness, Shannon-Wiener diversity, and Pielou’s evenness were analyzed as a three-way randomized block design with replication (sites treated as blocks nested within stand development stage). Differences in Shannon-Wiener diversity index and ESV richness between decay classes was determined by multiple comparisons of analysis of variance using Tukey’s honestly significant difference (HSD) test with a threshold of p<0.05.

We used multivariate regression trees (MRTs) on rarefied, presence/absence data to compare assemblages of fungal ESVs across stand development stage, decay class, and tree species using the “mvpart” package in R (De’ath, 2002; De’ath, 2011). The final model was chosen based on a 1000 cross-validation process, picking the smallest tree that fell within one standard deviation of error.

A sample x ESV table containing read counts was converted to a presence-absence matrix. Binary Bray-Curtis dissimilarities were calculated using the sample x ESV table with the function *vegdist*. A Principal Coordinate Analysis (PCoA) was run using Bray-Curtis dissimilarities with the function *pcoa* from the package “ape” (Paradis and Schliep, 2019). PCoA scores were extracted from the first 2 axes and retained for Akaike Information Criterion (AICc) modelling (n=59). Coarse woody debris physical and chemical properties were fitted to the ordination plot using the function *envfit*.

The sample x ESV table was also used with the *vegdist* function to calculate binary Bray-Curtis dissimilarities. Beta dispersion was calculated for ESVs across stand development stage, site (within stand development stage), decay class, and wood type using the *betadisper* function, and heterogeneity of beta dispersions within groups was checked. Heterogenous beta dispersions were identified for decay class only. Permutational multivariate analysis of variance (PERMANOVA) using the binary Bray-Curtis dissimilarity matrices was run using the function *adonis* to test the significance of groupings (stand development stage, site (within stand development stage), decay class, tree species).

The relationship of wood physical and chemical properties and carbon mineralization with fungal community composition was assessed using backwards-stepwise second-order Akaike Information Criterion (AICc) modelling using the function *dredge* from the package “MuMIn” (Barton, 2019). AICc is recommended when n/k < 40, where k is the number of parameters and n is the sample size (Burnham & Anderson, 2002). In this case, k=12 and n=59 (<40), therefore AICc was utilized. Pearson’s correlation coefficients were calculated using the function *rcorr* from the package “MASS” (Venables & Ripley, 2002). A cut-off of r = 0.7 was used to determine multicollinearity (Dormann et al., 2013). Density (g cm^-3^), total nitrogen (%), calcium (mg kg^-1^), and magnesium (mg kg^-1^) had r>0.7 and were removed prior to AICc modelling (Table S3). These variables were chosen for removal in order to retain correlated variables of ecological significance in the models (e.g., C/N ratio and P retained over N).

**Table 3:** Diversity metrics of fungal communities within coarse woody debris across stand development stage, tree species, and decay class (standard deviation in parentheses). Values within columns sharing the same letter do not differ significantly as determined by multiple comparisons using Tukey’s HSD test (*p*<0.05). Diversity metrics completed on 59 pooled CWD pieces spanning 11,178 ESVs.

|  |  | ESV richness (S) | Pielou's Evenness (J) | Shannon-Wiener Diversity (H) |
| --- | --- | --- | --- | --- |
| Decay Class | 1 | 248<sup>a</sup> (156) | 0.53 (0.15) | 2.89<sup>a</sup> (0.93) |
|  | 2 | 257<sup>a</sup> (92) | 0.6 (0.09) | 3.31<sup>b</sup> (0.66) |
|  | 3 | 541<sup>b</sup> (164) | 0.6 (0.09) | 3.77<sup>bc</sup> (0.74) |
|  | 4 | 563<sup>b</sup> (139) | 0.65 (0.05) | 4.11<sup>c</sup> (0.43) |
|  | 5 | 535<sup>b</sup> (231) | 0.61 (0.08) | 3.79<sup>bc</sup> (0.69) |
| Tree Species | Jack pine | 429 (220) | 0.59 (0.14) | 3.53 (0.99) |
|  | Black spruce | 456 (456) | 0.58 (0.09) | 3.53 (0.67) |
|  | Trembling aspen | 453 (239) | 0.62 (0.07) | 3.72 (0.7) |
| Stand Development Stage | Self-thinning | 409 (220) | 0.6 (0.12) | 3.55 (0.9) |
|  | Mature | 482 (206) | 0.6 (0.08) | 3.68 (0.67) |

## 3. Results

### 3.1 Chemical and physical properties, and C mineralization rates of CWD

Site by site comparisons of the same tree species and decay stage in self-thinning and mature stands were generally not explored in detail as part of this study as C mineralization rates (p=0.37), density (p=0.55), total carbon (p=0.59), total nitrogen (p=0.93), C/N ratio (p=0.67), N/P ratio (p=0.69), phosphorous (p=0.23) and potassium (p=0.10) concentrations did not significantly differ between stand development stages (Table 2; Table S4). However, we note that diameter (p=0.049), as well as calcium (p=0.035), magnesium (p=0.0012), and manganese (p=7.4e^-6^) concentrations in CWD, were significantly higher, and moisture content (p=3.6e^-4^) was significantly lower, in the mature stands (Table 2).

**Table 4:** Individual fungal ESV variances by split within the multivariate regression tree (MRT), total variance of the tree, and total variance explained for the MRT.

| Fungal<br>ESVs | Split 1 | Split<br>2 | Split<br>3 | Split<br>4 | Split<br>5 | Split<br>6 | Split<br>7 | Split<br>8 | Split<br>9 | Split 10 | Split<br>11 | Split<br>12 | Split<br>13 | Split<br>14 | Split<br>15 | Split 16 | Tree Total | ESV Total |
| --- | --- | --- | --- | --- | --- | --- | --- | --- | --- | --- | --- | --- | --- | --- | --- | --- | --- | --- |
| fITS94R<br>_Otu1 | 1.9e<sup>-3</sup> | 4.8e<sup>-3</sup> | 2.9e<sup>-3</sup> | 3.9e<sup>-3</sup> | 2.0e<sup>-4</sup> | 0 | 8.8e<sup>-3</sup> | 0 | 0 | 3.0e<sup>-4</sup> | 0 | 0 | 6.0e<sup>-4</sup> | 1.3e<sup>-3</sup> | 0 | 0 | 0.025 | 0.042 |
| fITS94R<br>_Otu10 | 2.1e<sup>-3</sup> | 2.1e<sup>-3</sup> | 3.0e<sup>-4</sup> | 0 | 1.7e<sup>-3</sup> | 0 | 2.0e<sup>-4</sup> | 4.0e<sup>-4</sup> | 7.0e<sup>-4</sup> | 6.0e<sup>-4</sup> | 0 | 0 | 6.0e<sup>-4</sup> | 1.0e<sup>-4</sup> | 0 | 0 | 8.9 e<sup>-3</sup> | 0.030 |
| fITS94R<br>_Otu11 | 2.1e<sup>-3</sup> | 0 | 1.0e<sup>-4</sup> | 1.0e<sup>-3</sup> | 2.1e<sup>-3</sup> | 2.0e<sup>-4</sup> | 2.2e<sup>-3</sup> | 4.0e<sup>-4</sup> | 0 | 1.8e<sup>-3</sup> | 0 | 1.0e<sup>-3</sup> | 6.0e<sup>-4</sup> | 1.3e<sup>-3</sup> | 0 | 0 | 0.013 | 0.030 |
| fITS94R<br>_Otu13 | 2.1e<sup>-3</sup> | 5.2e<sup>-3</sup> | 3.0e<sup>-4</sup> | 2.2e<sup>-3</sup> | 0 | 0 | 1.0e<sup>-3</sup> | 1.0e<sup>-4</sup> | 0 | 0 | 0 | 0 | 1.0e<sup>-4</sup> | 0 | 0 | 0 | 0.011 | 0.030 |
| fITS94R<br>_Otu14 | 0.018 | 0 | 4.0e<sup>-3</sup> | 2.0e<sup>-4</sup> | 2.0e<sup>-4</sup> | 2.0e<sup>-4</sup> | 2.0e<sup>-4</sup> | 0 | 0 | 3.0e<sup>-4</sup> | 2.0e<sup>-4</sup> | 0 | 0 | 1.3e<sup>-3</sup> | 7.0e<sup>-4</sup> | 2.9e<sup>-3</sup> | 0.029 | 0.055 |
| fITS94R<br>_Otu2 | 3.0e<sup>-3</sup> | 9.0e<sup>-4</sup> | 8.1e<sup>-3</sup> | 3.9e<sup>-3</sup> | 1.0e<sup>-4</sup> | 1.0e<sup>-3</sup> | 3.9e<sup>-3</sup> | 2.0e<sup>-4</sup> | 0 | 1.0e<sup>-4</sup> | 0 | 2.0e<sup>-4</sup> | 5.3e<sup>-3</sup> | 6.0e<sup>-4</sup> | 0 | 7.0e<sup>-4</sup> | 0.028 | 0.065 |
| fITS94R<br>_Otu4 | 1.2e<sup>-3</sup> | 3.5e<sup>-3</sup> | 7.0e<sup>-4</sup> | 2.0e<sup>-4</sup> | 2.0e<sup>-4</sup> | 2.0e<sup>-4</sup> | 1.0e<sup>-3</sup> | 9.0e<sup>-4</sup> | 0 | 9.0e<sup>-4</sup> | 2.0e<sup>-4</sup> | 1.0e<sup>-3</sup> | 6.0e<sup>-4</sup> | 0 | 7.0e<sup>-4</sup> | 0 | 0.012 | 0.039 |
| fITS94R<br>_Otu45 | 9.0e<sup>-4</sup> | 2.7e<sup>-3</sup> | 1.0e<sup>-4</sup> | 2.2e<sup>-3</sup> | 0 | 2.0e<sup>-4</sup> | 1.0e<sup>-3</sup> | 1.0e<sup>-4</sup> | 0 | 0 | 0 | 1.0e<sup>-3</sup> | 2.3e<sup>-3</sup> | 0 | 0 | 0 | 0.011 | 0.024 |
| fITS94R<br>_Otu66 | 2.0e<sup>-4</sup> | 1.7e<sup>-3</sup> | 1.0e<sup>-4</sup> | 1.0e<sup>-3</sup> | 0 | 2.0e<sup>-4</sup> | 1.0e<sup>-3</sup> | 9.0e<sup>-4</sup> | 0 | 0 | 0 | 1.0e<sup>-3</sup> | 6.0e<sup>-4</sup> | 0 | 0 | 0 | 6.6e<sup>-3</sup> | 0.024 |
| fITS94R<br>_Otu7 | 9.0e<sup>-4</sup> | 4.0e<sup>-4</sup> | 3.0e<sup>-4</sup> | 0 | 1.0e<sup>-4</sup> | 0 | 1.0e<sup>-3</sup> | 4.0e<sup>-4</sup> | 7.0e<sup>-4</sup> | 1.0e<sup>-4</sup> | 0 | 0 | 6.0e<sup>-4</sup> | 6.0e<sup>-4</sup> | 0 | 0 | 5.1e<sup>-3</sup> | 0.024 |
| Omitted<br>ESVs<br>(11,168) | 5.27 | 2.96 | 2.65 | 2.28 | 2.38 | 2.31 | 1.59 | 1.88 | 1.65 | 2.07 | 2.41 | 2.26 | 1.37 | 1.79 | 1.46 | 1.81 | 36.1 | 99.6 |
| Total | 5.30 | 2.98 | 2.67 | 2.30 | 2.39 | 2.31 | 1.61 | 1.89 | 1.64 | 2.08 | 2.41 | 2.26 | 1.38 | 1.79 | 1.46 | 1.81 | 36.3 | 100 |

Across CWD species, there were variations in physical, chemical, and biological characteristics of CWD (Table 2). Overall (all decay classes combined), trembling aspen CWD had significantly lower density (p=0.001), total carbon (p=9.9e^-7^), C/N ratios (p=1.3e^- 13^), and manganese concentrations (p<2.2e^-16^) than black spruce and jack pine (Table 2; Table S4). Trembling aspen CWD had significantly higher C mineralization than black spruce and jack pine CWD (p=0.0043), as well as moisture contents (p=2.4e^-4^), total nitrogen (p=2.5e^-12^), and phosphorous (p=7.3e^-11^), potassium (p=1.3e^-12^), calcium (p<2.2e^- 16^), and magnesium (p<2.2e^-16^) concentrations (Table 2; Table S4).

Across decay classes there were significant variations in all CWD physical and chemical wood properties, except for total carbon (p= 0.32) and magnesium (p=0.65) concentrations (Table 2; Table S4). Wood density (p<2.2e^-16^) significantly decreased with decay class, which contrasted with increasing moisture content (p<2.2e^-16^) (Table 2). Both total nitrogen (p<2.2e^-16^) and phosphorous (p<2.2e^-16^) concentrations remained stable across decay classes 1-3 before increasing in decay classes 4 and 5 (Table 2). Potassium concentrations (p=0.027) were highest in decay class 1, significantly decreased in decay class 2, and then remained stable across the more advanced (DC3-5) decay classes (Table 2). Both calcium (p=0.011) and manganese (p=4.8e^-4^) concentrations remained stable across decay classes 1, 2 and 3 before significantly increasing in decay class 4 (Table 2). C/N ratio (p<2.2e^-16^) remained stable across decay classes 1 and 2 before significantly decreasing across decay classes 3, 4 and 5 (Table 2). There were significant increases in N/P ratio (p=0.0077) from decay class 1 to decay class 2 and 3 before decreasing in decay classes 4 and 5 (Table 2). In terms of an index of biological activity, C mineralization (p=9.0e^-9^) significantly increased with decay class, with the highest rates being detected in the decay class 5 wood sections.

### 3.2 Alpha Diversity

When looking at alpha diversity of fungal communities across CWD pieces, there were no significant (p<0.05) differences in Shannon-Wiener diversity or ESV richness across stand development stage, sites (within stand development stage), or tree species (Table 3; Table S5). There were also no significant (p<0.05) differences in Pielou’s evenness across stand development stage, sites (within stand development stage), tree species, or decay class (Table 4; Table S5). Shannon-Wiener diversity (p=0.0036) and ESV richness (p=2.3e^-5^) did significantly increase with decay class; with both Shannon-Wiener diversity and ESV richness highest in decay class 4 with values of 4.11 and 563, respectively (Table 3; Table S5). However, Shannon-Wiener diversity and ESV richness in decay class 4 were only significantly different from decay classes 1 and 2, suggesting more diverse fungal communities occur in intermediate to late decay stages (Table 3).

**Table 5:**
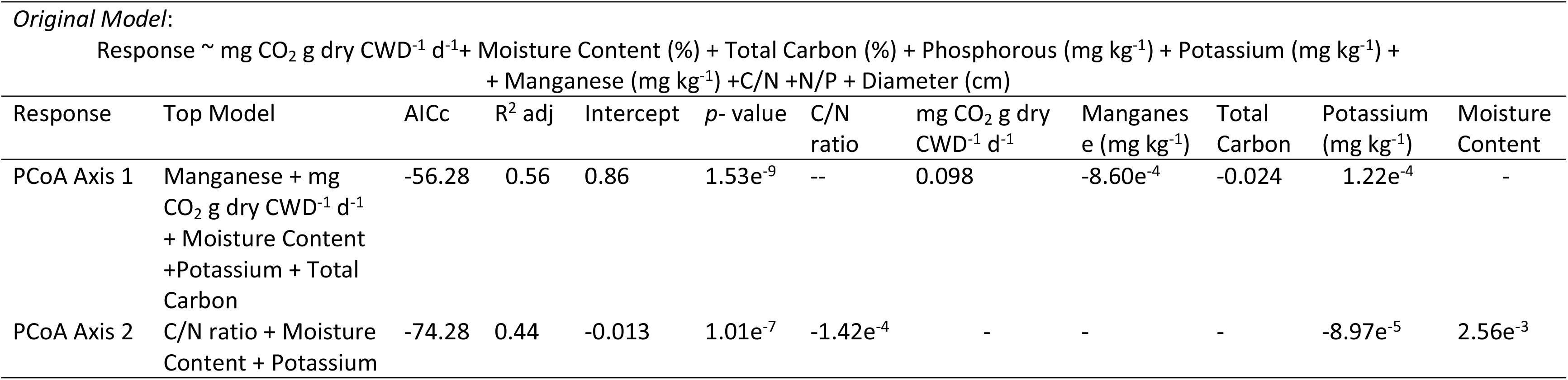
Top scoring backwards-stepwise second-order Akaike Information Criterion (AICc) model predictor variables and results with Principal Coordinate Analysis (PCoA) scores (representing fungal communities across tree species and decay class) as the response variables.

### 3.3 Beta Diversity

The total variance explained by the fungal presence/absence MRT was 36.3% (Table 4). We found that the primary determinant factor in differentiating fungal community composition in CWD was tree species, with 5.30% of total variance explained by the first split of conifer (black spruce and jack pine) versus hardwood (trembling aspen) (Figure 2; Table 4). For both conifers (split 2 explaining 2.98% of the total variance) and trembling aspen (split 3 explaining 2.67% of the total variance), the next most influential factor in determining fungal communities was decay stage, with the initial decay stages (decay class 1-3 in conifers and decay class 1-2 in aspen) separated from later decay stages (Figure 2; Table 4). The third most influential split for both conifers (split 4 explaining 2.30% of the total variance) and hardwoods (split 6 explaining 2.31% of the total variance) was stand development stage (separating self-thinning and mature stands), perhaps due to changes in community from initial colonizer fungi (early decay stages in self-thinning stands) to more specialized wood decaying fungal species (late decay stages and mature stands) (Figure 2).

**Figure 2:**
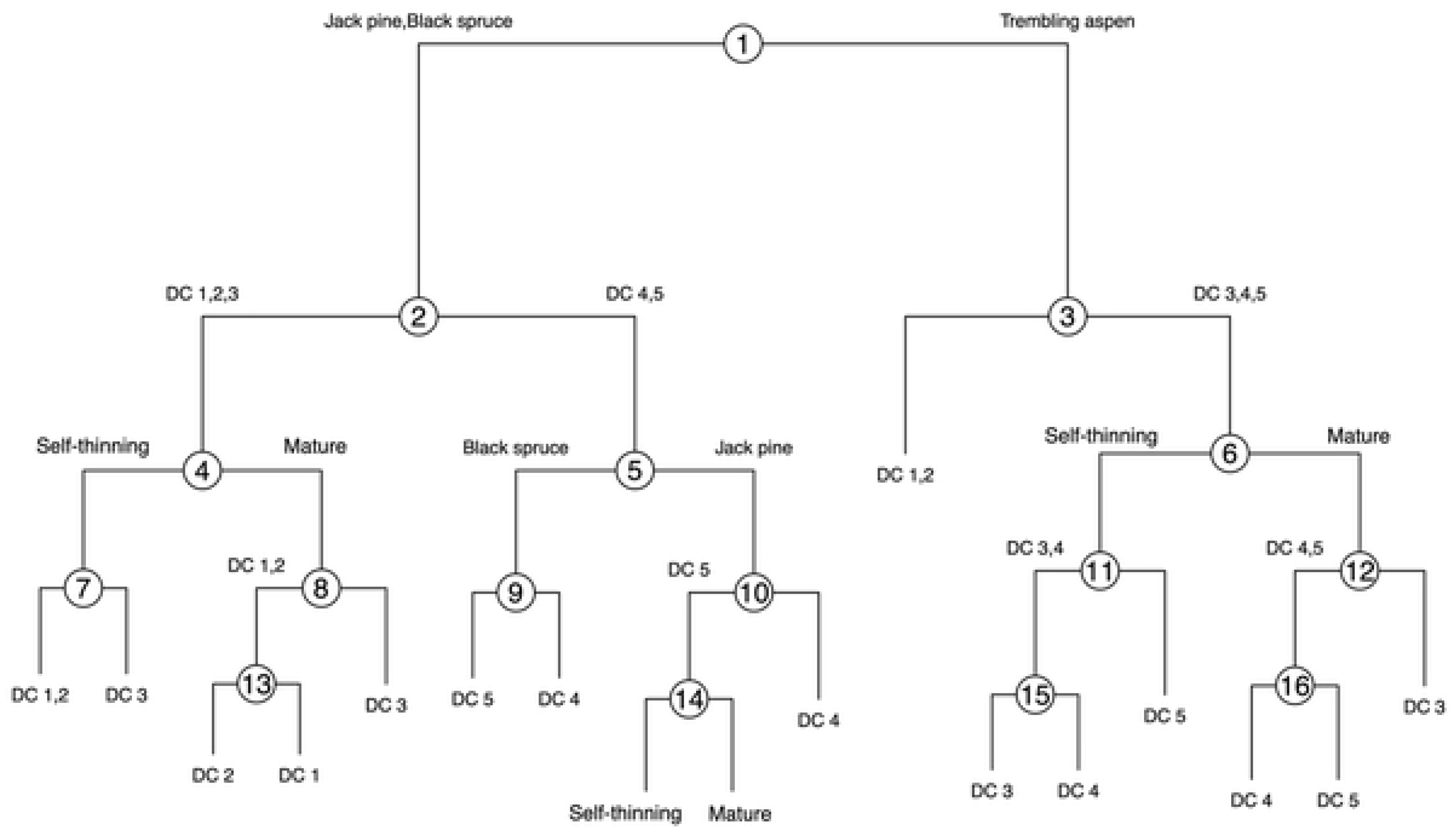
Multivariate regression tree (MRT) for fungal ESVs across stand development stage (self-thinning and mature stands), tree species (jack pine, black spruce, trembling aspen), and decay class (DC 1, DC 2, DC 3, DC 4, DC 5).

Continuing down the left-hand side of the MRT, early decay stage (DC1-2) conifers in both self-thinning and mature stands were split from decay class 3 (splits 7 and 8 explaining 1.67% and 1.87% of the total variance, respectively), perhaps signifying the start of a transition to a more specialized community of wood decay fungi (Figure 2; Table 4). Decay classes 4 and 5 were split by CWD species, separating black spruce and jack pine (split 5 explaining 2.39% of the total variance), suggesting that for later decay stage conifer CWD, species is more influential than stand development stage (Figure 2; Table 4). Black spruce CWD was further split between decay class 4 and 5 (split 9 explaining 1.66% of the total variance) (Figure 2; Table 4). Jack pine CWD was also split further, separating decay class 4 and 5, however, it also separated decay class 5 CWD by stand development stage (splits 10 and 14 explaining 2.08% and 1.79% of the total variance, respectively) (Figure 2; Table 4).

The right-hand side of the MRT dealt with splits associated with the hardwood species (trembling aspen) CWD (Figure 2). For trembling aspen CWD, decay classes 1 and 2 were separated from decay classes 3-5 (split 3 explaining 2.67% of the total variance) (Figure 1; Table 4). This split suggests that for hardwood CWD decay class 3s, the fungal communities may be more similar to late-stage communities compared to early, colonizing communities. Decay classes 3, 4, and 5 were further split by stand development stage (split 6 explaining 2.31% of the total variance) (Figure 2; Table 4). In the self-thinning stands, decay classes 3 and 4 were split from decay class 5 (split 11 explaining 2.41% of the total variance) (Figure 2; Table 4). In the older stands (mature to over-mature), decay classes 4 and 5 were split from decay class 3 (split 12 explaining 2.26% of the total variance) (Figure 1; Table 4). From the various splits by decay class in late-stage hardwood and conifer CWD, results would suggest that fungi are undergoing dynamic changes in community composition as decay progresses.

Supporting the results from the MRT, the PCoA analysis also showed separations of fungal communities along axis 1 between all three tree species sampled (trembling aspen, jack pine, or black spruce (Figure 3a). The PERMANOVA also showed that species composition did significantly differ between tree species (p=0.001) (Table S6). The ordination analysis showed graduated separation of fungal communities along axis 2 as wood decay advanced through the five decay classes, and the PERMANOVA showed that species composition was significantly different among decay classes (p=0.001) (Figure 3b; Table S6). Although there is considerable overlap between the stand development stages along the axis 1, fungal communities in the stands in their self-thinning stage of stand development did tend to separate along axis 2 (Figure 3c). The PERMANOVA analysis showed that species composition significantly differed between stand development stage (p=0.001) (Table S7). We found that species composition differed significantly among sites (p=0.011); however, when accounting for stand development stage, site becomes insignificant (p=0.134) (Table S6; Table S7). This suggests that stand development stage, not individual site differences, is, at least, partially controlling difference in fungal community composition.

**Figure 3:**
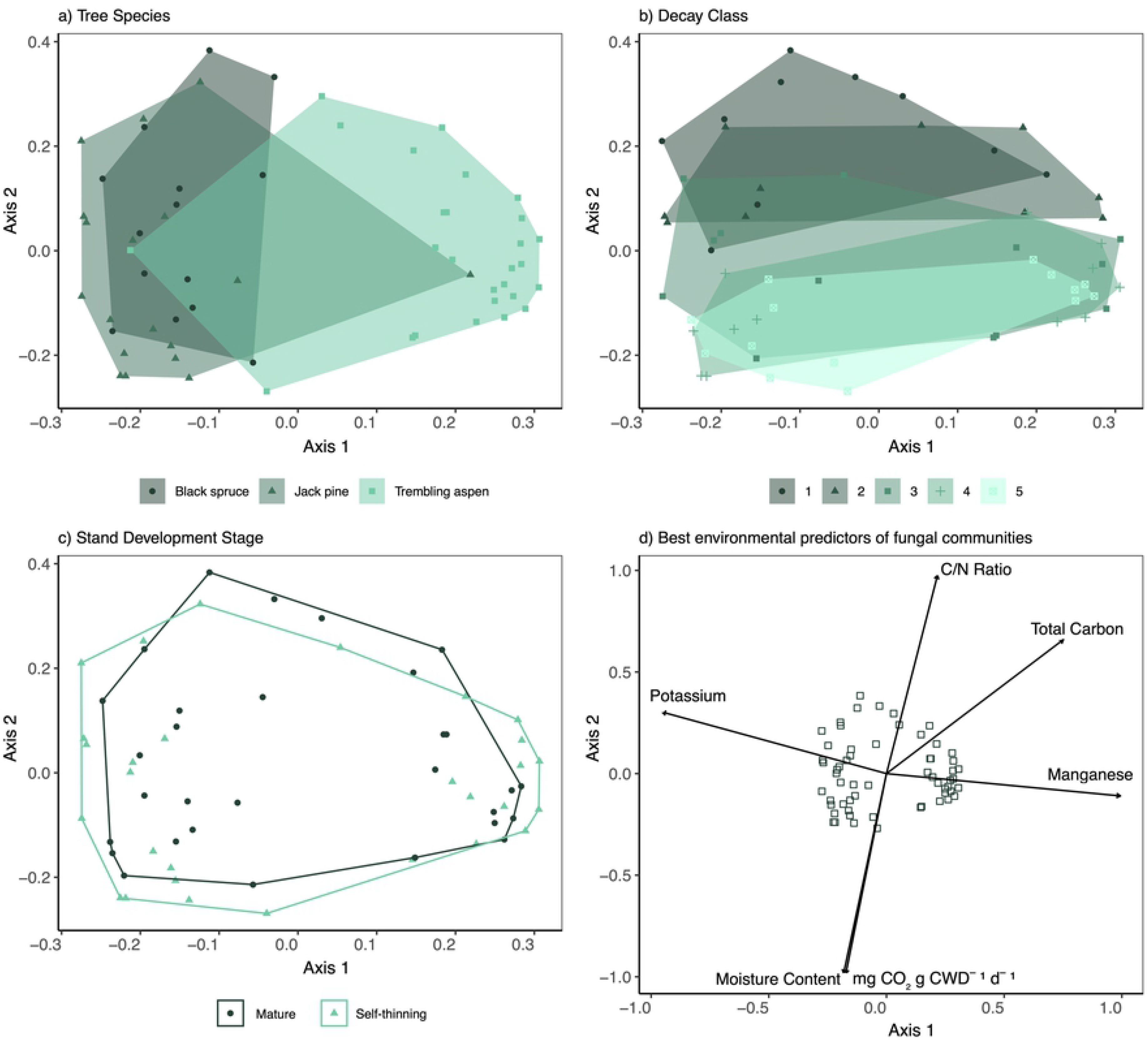
Principal Coordinate Analysis (PCoA) plots of fungal communities across stand development stage, tree species, and decay class, and with associated environmental predictors. Diversity metrics completed on 59 pooled CWD pieces spanning 11,178 ESVs. [Print in colour]

Additionally, we found there was a significant interaction between tree species and decay class (p=0.005) (Table S6).

### 3.4 Examining the relationships between fungal community shifts and changes in the physical, chemical, and biological properties of CWD

To investigate the relationship between fungal communities and the physical, chemical, and biological properties of CWD, we used AICc model selection with scores extracted from the first 2 axes of the PCoA plot representing fungal communities across tree species and decay class as the response variables, and C mineralization (mg CO^2^ g CWD^-1^ day^-1^), moisture content (%), total carbon (%), diameter (cm), concentrations (mg kg^-1^) of phosphorous, potassium, magnesium, and manganese, and C/N and N/P ratios as the predictor variables (n=59) (Table 5). With our PCoA axis 1 as the response variable representing fungal community composition across CWD tree species, we identified Mn and K concentrations (mg kg^-1^), total carbon (%), moisture content (%) and mg CO_2_ g dry CWD^-1^ d^-1^ as the best predictors of community composition across CWD tree species (Table 5). Additionally, we used AICc model selection with our PCoA axis 2 as the response variable representing fungal community composition across CWD decay class and identified K concentration (mg kg^-1^), C/N ratio, and moisture content (%) as the best predictors of community composition across decay class (Table 5). Mn, and K concentrations, C/N ratio, total carbon (%), moisture content (%) and mg CO_2_ g dry CWD^-1^ d^-1^ had p values <0.05 when fitted to the ordination plot (Figure 2d). Unidirectional causal relationships could not be established, as the interplay of initial CWD properties influencing fungal communities that themselves modify wood properties through decay stages are highly dynamic and interactive.

### 3.5 Taxonomic and Functional Fungal Composition

There was a total of 11, 178 ESVs identified across samples (n=129). Of the 11,178 ESVs, 4,911 were identified to class with at least 75% accuracy.

All ESVs (11,178) were run against the FUNGuild database to determine traits. ESVs were classified to trait with a possible, probable, and highly probable confidence ranking. Of the 11,178 ESVs run against the FUNGuild database, 6,882 were unassigned, and 3,372 could not be classified to trait. As a result, only 924 ESVs were assigned to a trait (white rot fungi, brown rot fungi, brown rot-white rot fungi, soft rot fungi, hypogeous fungi, poisonous fungi, and blue-staining fungi). FUNGuild results may not be representative of the entire fungal community in CWD but are more a reflection of what we presently know about fungal traits, as current databases are far from complete.

In terms of ESVs identified to class, Agaricomycetes, Leotiomycetes, and Eurotiomycetes were the dominant classes across all CWD species, likely due to the broad range of wood pathogens and wood decay fungi, including brown, white, and soft rot, within these classes (Figure 4). We can see these classes identified to trait when looking at the ESVs classified by FUNGuild: trembling aspen CWD had higher proportions of white rot fungi compared to conifer CWD, and conifer CWD had higher proportions of brown rot fungi (Figure 5). Trembling aspen CWD had higher proportions of Sordariomycetes and Dothideomycetes compared to conifer CWD, which we saw reflected in the higher proportions of blue-staining and poisonous fungi (as classified by FUNGuild) (Figure 4, Figure 5). Sordariomycetes and Dothideomycetes both include several genera of wood pathogens, including blue-staining fungi. ESVs classified as “poisonous” by FUNGuild, are primarily *Gyromitra infula* (class Pezizomycetes) and *Psilocbye montan*a (class Agaricomycetes), both of which are commonly located in coniferous forests throughout North America. Black spruce CWD contained higher proportions of brown rot-white rot fungi, compared to jack pine or trembling aspen (Figure 5). FUNGuild identified “brown rot-white rot” fungi as primarily being from the family Polyporaceae, which contains both brown rot and white rot fungi (Figure 5).

**Figure 4:**
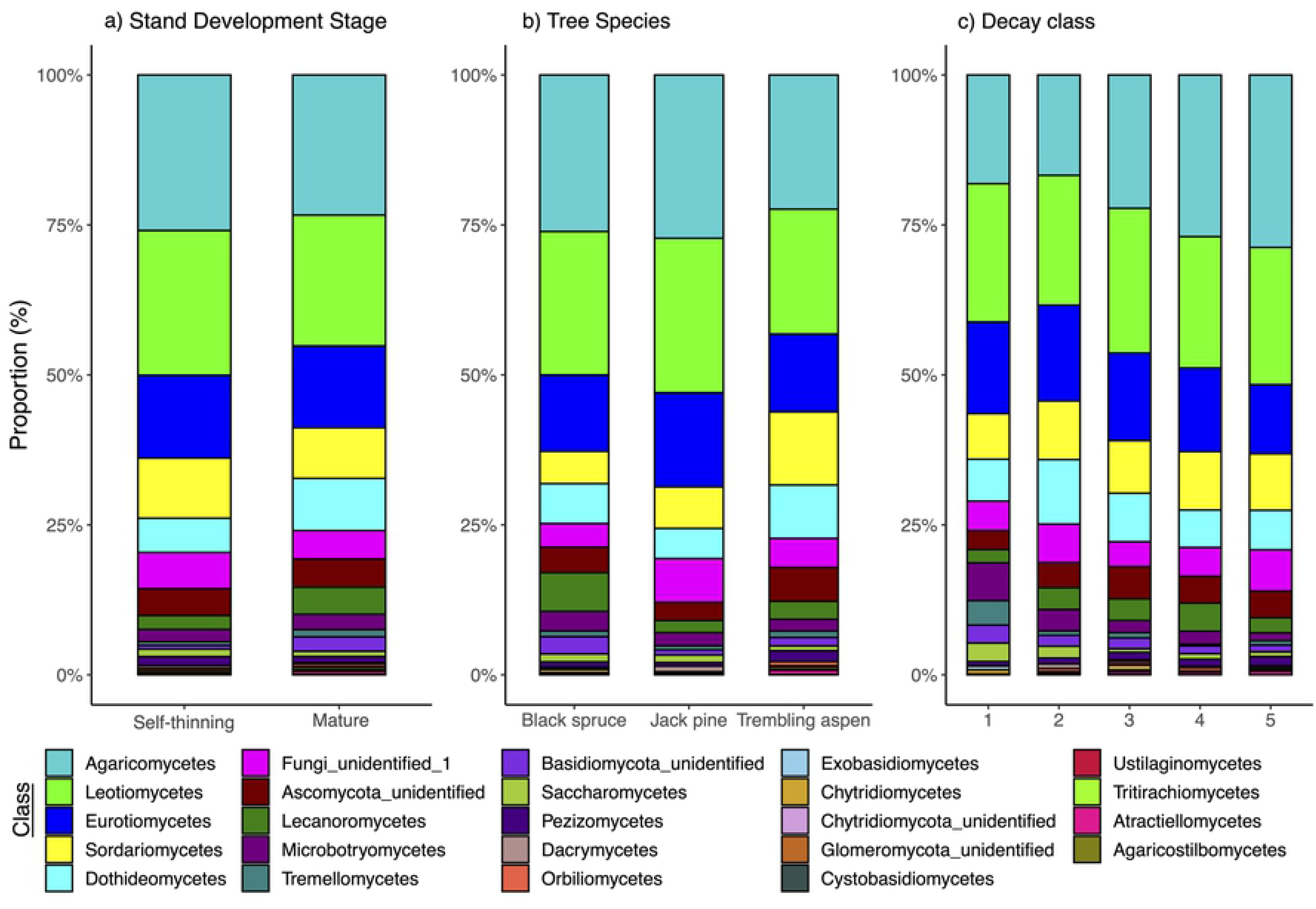
Proportion of total fungal ESVs across stand development stage, tree species, and decay class, classified to Class with 50% cut-off for fragments <250bp across 59 pooled CWD pieces and 4,911 ESVs. [Print in colour]

**Figure 5:**
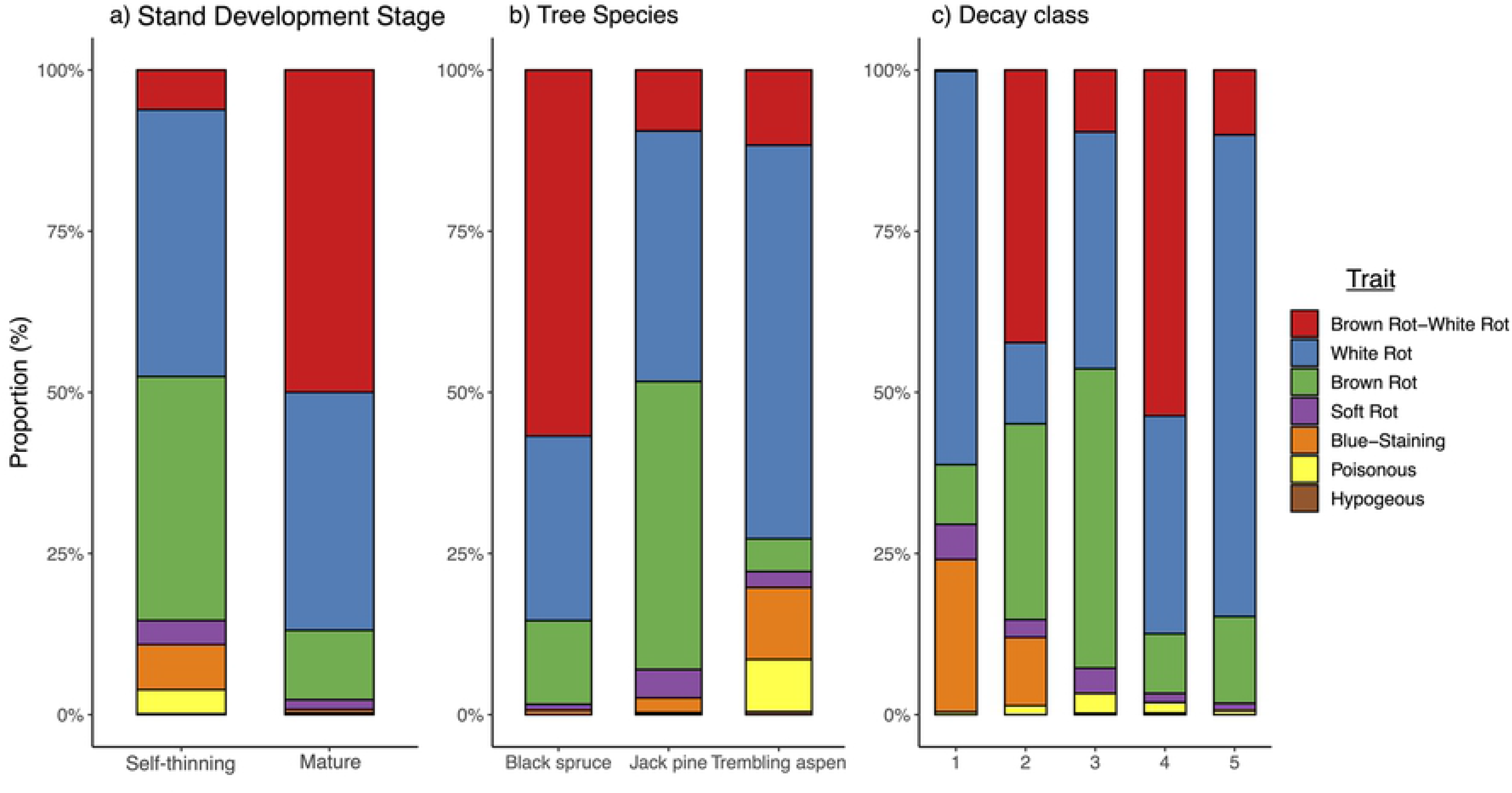
Proportion of ESVs that were assigned a trait across stand development stage, tree species, and decay class, classified to functional trait with possible, probable, and highly probable confidence rankings across 59 pooled CWD pieces and 924 ESVs. [Print in colour]

Across decay class, we saw a shift toward increased proportions of Agaricomycetes and decreased proportions of Leotiomycetes and Eurotiomycetes as wood decay proceeds (Figure 4). As these classes all contain species of wood decay fungi amongst other types of fungi (wood pathogens, mycorrhizal), this shift is likely occurring at a species level, with most of the wood decay fungi species present in the later decay stages belonging to Agaricomycetes. We saw this reflected in the ESVs classified to trait by FUNGuild: although all five decay classes had some proportion of white rot, brown rot, and soft rot fungi, we saw increased proportions of white rot fungi and brown rot fungi in decay class 5 compared to decay class 1 (Figure 5). Additionally, we saw increased proportions of blue-staining and poisonous fungi (Sordariomycetes, and Pezizomycetes and Agaricomycetes, respectively) in decay classes 1, 2, and 3, perhaps signifying the colonization of CWD by wood pathogens that facilitate decay (Figure 5).

We found that Agaricomycetes was the dominant class across both stand development stages, likely due to Agaricomycetes containing many common species of fungi specialized for wood decay, both white rot and brown rot fungi (Figure 4). Leotiomycetes and Eurotiomycetes, both containing mycorrhizae and/or fungal parasites, and wood decay fungi, were also dominant across both stand development stages (Figure 4). The self-thinning stands had higher proportions of Sordariomycetes, whereas the mature stands had higher proportions of Dothideomycetes (Figure 4). Looking at ESVs classified to functional trait via FUNGuild, we saw high proportions of white rot and brown rot fungi (likely corresponding to Agaricomycetes, Leotiomycetes) in both the self-thinning and mature stands (Figure 5). The self-thinning stands also had higher proportions of soft rot fungi, corresponding to Sordariomycetes (Figure 5). In contrast, the mature stands had higher proportions of brown rot-white rot fungi, in this case Polyporaceae (Figure 5). Finally, the self-thinning stands had higher proportions of blue-staining fungi, all of which were identified to belong to the class Sordariomycetes (Figure 4, Figure 5).

## 4. Discussion

Our objective in this study was to examine fungal community structure within CWD, focusing on two key questions: *how does the fungal community composition in CWD change across stand development stage (stand age), tree species, and decay class, and what are the relationships between changes in fungal communities and the measured differences in the physical, chemical, and biological properties of CWD?*

### 4.1 Differences in fungal community composition in CWD as a function of stand development stage, tree species, and decay class

We found that CWD tree species was the primary factor influencing fungal community composition, which is consistent with previous Northern European studies that found significant differences in fungal community composition between European beech and Norway spruce, likely related to the structural composition of wood (e.g., proportions of cellulose, hemicellulose, lignin) and the differing biochemical systems associated with fungal species responsible for degrading these components (Baber et al., 2016; Hoppe et al., 2016; Schwarze, 2007). More specifically, the ratio of cellulose, hemicellulose, and lignin in CWD varies between conifer and hardwood species, with hardwoods generally having lower lignin, equal cellulose, and higher hemicellulose contents (Harmon et al., 2004). White rot fungi are commonly associated with hardwood CWD (Schwarze, 2007), as confirmed by our findings. The rationale for this preference remains unclear but may be due to the type of lignin present in hardwood trees: hardwood trees contain equal parts guaiacyl and syringyl lignins, the latter of which has been shown to degrade more rapidly (Kirk et al., 1975). This difference would allow white rot fungi to degrade hardwood CWD easier and faster than conifer CWD, as conifers primarily have guaiacyl lignin present (Schwarze, 2007). In contrast, black spruce and jack pine CWD had a higher proportion of brown rot fungi, with the literature commonly associating brown rot fungi with conifer CWD (Schwarze, 2007). Similarly, the rationale for this the preference of fungi for wood type is unknown but could be related to the structural differences between conifers and hardwoods, particularly in parenchyma cells: in a study examining the resistance of parenchyma cells in wood to degradation by brown rot fungi, Schwarze et al. (2003) found that weight loss associated with brown rot fungi wood degradation was correlated with parenchyma content, with higher weight loss and lower parenchyma content in Norway spruce wood blocks.

We found that decay stage was the next most influential factor in determining fungal community composition. In particular, late decay stage communities differed significantly from early colonizing fungal communities as decayed advanced, with the proportions of specialized wood decay fungi increasing through decay classes, which is consistent with previous literature (Rajala et al., 2010). Fungal diversity and richness have been shown to increase with decay class, beginning with pioneer, colonizing fungi (Hoppe et al., 2016; Kubartová et al., 2012; Rajala et al., 2015). Fungal colonization of CWD ranges from spores of pathogenic, pioneer fungi already present in tree trunks before they fall, to spores entering through the cut-ends of felled trees, to spores colonizing CWD via air, water currents, rain splash, animal vectors, to arriving as vegetative mycelium (Boddy, 2001; Fukasawa et al., 2009). We found higher proportions of blue-staining fungi in early decay classes compared to intermediate and late decay classes: blue-staining fungi are considered early colonizers as they infect living trees via hyphal growth in sapwood, eventually causing tree death (Ballard & Walsh, 1984).

Fungal communities within CWD are continually changing both temporally and spatially as wood decomposes, contrary to the notion of a climax community at any particular stage of decay (Boddy, 2001). Our results suggest that as CWD decays, the influence of colonizer fungi decreases and stabilized fungal communities form (Rajala et al., 2010). After CWD has been colonized, community development is influenced by interspecific interactions and species-specific substrate modification (Boddy, 2001; Kubartová et al., 2012). Kubartová et al. (2012) found that when many species coexist the indirect effects of species interactions are likely positive and may alleviate the negative effects of competition seen with increasing diversity and less available space. As CWD becomes more decayed, the quality of the substrate and remaining consumable energy decreases and more interacting fungi occupy space within CWD, selecting for fungi that have complimentary functional traits (Kubartová et al., 2012). For example, Agaricomycetes were the most abundant in decay class 5, likely due to their ability to select for fungi with complementary fungal traits, including production of secondary metabolites (e.g., Botales sp.), forming hyphal mats, and mycoparasitism, as wood becomes more decayed (Hibbett et al. 2014). The class Agaricomycetes is comprised of a diverse array of 16,000 species that decay wood, act as pathogens and parasites, and are lichen-forming or ectomycorrhizal (Kohler et al., 2015; O’Hanlon & Harrington, 2011).

The least influential factor in determining fungal community composition was stand development stage (stand age classes). Decreases in fungal species richness and diversity in CWD from managed silvicultural stands compared to natural, mature stands is well noted in the literature (Bader et al., 1995; Lõhmus, 2011; Müller et al., 2007; Penttilä et al., 2004). While we did not see significant differences in ESV richness or diversity between fungal communities in CWD between self-thinning and mature stands, we did see a clear separation of fungal communities in our ordination analysis between stand development stages. These results are consistent with studies examining fungal community composition across stands with different ages and/or successional stages in northern Europe (Junninen et al., 2006; Kubart et al., 2016). Fungi in CWD have been shown to prefer closed canopies, likely due to reduced sunlight and higher humidity and ground surface moisture compared to the conditions in open canopy stands, that, in turn, influence how communities form in forests as they development (Horák et al., 2016). Odriozola et al. (2020) found that community composition of soil fungi in mixed temperature forests in the Czech Republic changed with stand age, likely due to shifts in nutrient supply from plant/tree hosts to root fungi, as well as changes in soil chemistry and ground vegetation. Though we did not examine soil fungi in this study, they are often in close association with saprotrophic fungi by proximity.

### 4.2 Relationships between changes in fungal communities and the measured differences in the physical, chemical, and biological properties of CWD

We found that trembling aspen CWD had significantly higher rates microbial C mineralization (mg CO_2_ g dry CWD^-1^ d^-1^) than conifer CWD, consistent with previous literature examining CWD decomposition controls (Gough et al., 2007; Harmon et al., 2004; Herrmann & Bauhus, 2013; Kahl et al., 2015; Pastorelli et al., 2017; Wang et al., 2002). Herrmann & Bauhus (2013) incubated CWD *in situ* while controlling air temperature and humidity and found that CO_2_ flux was higher in European beech CWD compared to Norway spruce or Scots pine. Higher C mineralization in trembling aspen CWD may be, in part, due to the colonized fungal community: trembling aspen CWD are typically dominated by white rot fungal species that have the ability to decompose lignin as well as cellulose and hemicellulose; whereas black spruce and jack pine CWD are typically dominated by brown rot fungi that primarily decompose cellulose and hemicellulose (Schwarze, 2007).

We found that microbial C mineralization (mg CO_2_ g dry CWD^-1^ d^-1^) significantly increased with decay class, supported by Pastorelli et al. (2017), who found that CO_2_ production was significantly higher in later decay classes in European black pine (*Pinus nigra* Arnold) CWD in Central Italy, and Gough et al. (2007) who found that temperature-normalized respiration rates increased with decay class in both hardwood and conifer CWD from a north temperate forest in Michigan, USA. Furthermore, van der Wal et al. (2013) suggested that higher species richness enhances decomposition via synergistic interactions. Functional niche complementarity, also known as niche differentiation and facilitation, describes the hypothesis of increased performance of a community when compared to an individual species (Loreau et al., 2001). We found significantly higher Shannon-Wiener values in decay class 4, allowing us to infer that the more heterogeneous environment of later decay stage logs increased niche availability and resulted in positive interactions between fungal species. A more diverse community would have improved resource exploitation, and, in this case, result in increasing CO_2_ production (van der Wal et al., 2013). Mäkipää et al. (2017) found that species richness in black spruce CWD increased with decay class, eventually reaching the same species richness and community composition as the underlying soil. This increased species richness was partly due to the inclusion of mycorrhizal species from soil, which, when observed in conjunction with the increase in nitrogen and decrease in carbon in CWD, collectively can influence carbon dynamics and accelerate wood decay (Mäkipää et al., 2017).

Cations, including manganese and potassium, are required by fungi for growth, function, development, and reproduction (Jellison et al., 1997). We found that across CWD species, potassium and manganese were important predictors of fungal community composition, and across CWD decay class, potassium was an important predictor. This may be in part due to the chemical variation across tree species: hardwoods generally have higher nutrient contents than conifers (Harmon, 2004). Variations in cation concentrations in external environments (i.e., CWD) may influence community development, as fungi use cations in their biochemical systems to degrade wood and take up nutrients (Jellison et al., 1997).

Mn is an important regulator in lignin degradation of white rot fungi (Ostrofsky et al., 1997). White rot fungi breakdown lignin via lignin-modifying enzymes (LMEs) that include lignin peroxidase (LiP) (Bonnarme & Jeffries 1990). LiP is formed at low Mn concentrations and repressed at high Mn concentrations (Bonnarme & Jeffries, 1990). Low Mn concentration and high abundance of white rot fungi in trembling aspen CWD might suggest that LiP is being formed preferentially over other LMEs.

Variations in potassium concentrations across decay class may be caused by the presence/absence of different species as communities develop: Ostrofsky et al. (1997) found that K concentrations increased in wood inoculated with *Armillaria* sp. but decreased in wood inoculated with *T. versicolor*. Overall, they found most fungi do not accumulate K in CWD and suggest this is due to both K’s high solubility and cation-binding. In particular, Ca has been shown to preferentially accumulate instead of K in wood, and, when rainfall is a factor, K has been shown to leach at a faster rate than mass loss (Foster & Lang, 1982; Ostrofsky et al., 1997).

Conifer species generally have higher carbon content than hardwood species due to their higher lignin content, as lignin has the highest percentage of carbon of all wood macromolecules (Lamlom & Savidge, 2003). A higher abundance of brown rot fungi compared to white rot fungi in conifers may leave more carbon in the wood tissue, as brown rot fungi decompose slower than white rot fungi and do not decompose lignin (Jurgensen et al., 1989). Trembling aspen CWD in this study had a significantly higher concentration of N than conifer species, potentially due to the symbiotic relationship of trembling aspen with both ectomycorrhizal and arbuscular mycorrhizal fungi prior to tree mortality: mycorrhizal fungi are known to transfer N to their host plant which may influence fungal community composition after tree death (Brundrett et al., 1990; Vozzo & Hacskaylo, 1974). Both saprotrophic and mycorrhizal fungi have been found to sequester nitrogen (Leake et al., 2003).

There is a dynamic interplay between initial nutrient content of CWD and the microbial community driving nutrient input and export through the decay continuum, which influences the rate of decomposition (Harmon et al., 2004). Moisture content increases with decomposition and is inversely related to wood density (Harmon et al., 2004). We found that wood density significantly decreased, and moisture content and fungal diversity significantly increased as decay advanced, as confirmed by other studies (Kubartová et al., 2012; Rajala et al., 2012). As wood becomes more decomposed, its density decreases and moisture content increases, causing CWD to become more heterogeneous, increasing the availability of different niches and allowing for higher diversity (Boddy, 2001).

### 4.3 Management Implications

The results from this study can aid in the conservation of biodiversity and support the maintenance of fundamental ecosystem functions in boreal forests by helping inform forest policy, thus improving sustainable forest management strategies. For example, it has been commonly noted that the use of forest biomass as feedstock for the growing bio-economy sector can modify the supply of deadwood essential to forest ecosystem structure and function (Berch et al., 2011; Franklin, 1990; Hunt et al., 2010; Kerr, 1999; Moroni et al., 2015; Omari & Maclean, 2015; Puddister et al., 2011). Biomass harvesting in historically deadwood-dependent ecosystems has significantly decreased the amount and quality of deadwood present, which could lead to losses in habitat, reduce carbon storage, and affect a broad suite of ecosystem services (Venier et al., 2015). In some cases, deadwood reduction has been identified as a major contributor to reduced species diversity: in Sweden, deadwood loss is the primary threat to more than 60% of 739 threatened invertebrate forest species (Berg et al., 1994). Though Canadian biomass harvesting is less intensive than in Fennoscandia, the impacts on biodiversity and site productivity should continue to be taken into consideration in Canadian forest policy (Berch et al., 2011).

Forest policy across a number of jurisdictions applies general (coarse filter) and specific (fine filter) guidance to maintain deadwood quantities in support of the conservation of deadwood-dependent species (Berch et al., 2011; Bunnell & Houde, 2010; Titus et al., 2010). As a general rule, these guidelines commonly suggest minimizing the disturbance to pre-harvest deadwood, as well as retaining some number of live trees in harvested areas (i.e., green tree retention rule sets/thresholds). For example, in forest management guidelines in Ontario, Canada, live tree retention of at least 25 stems ha^-1^ is required in clearcut systems (OMNR, 2010). In addition to acting as immediate wildlife trees, these residual stems will eventually die or fall, adding to the deadwood pool over time. Beyond the recognized need for quantitative thresholds, our results also highlight that the qualitative characteristics of CWD (i.e., species, decay class distribution) must continue to be highly considered in the direction provided in forest management guidelines (Densmore, 2011; Fischer et al., 2012). For example, fungal community structure and successional dynamics could be altered with the preferential removal and/or retention of desirable tree species (e.g., conifer-based harvests), as we have described distinct fungal communities between conifer and hardwood CWD, and between early (decay class 1-2) and legacy (decay class 3-5) CWD. These results will assist in monitoring the effectiveness of existing forest management guidelines to provide a diversity of substrates that will help preserve a variety of species (Müller & Bütler, 2010; Tremblay et al. 2009; Work & Hibbert, 2011; Work et al., 2004).

## 5. Conclusion

Our work highlights the importance of maintaining a diverse pool of CWD, that consists of a variety of tree species and decay classes in managed forests. This direction would help to conserve forest biodiversity and preserve species dependent on coarse woody debris, but also support carbon sequestration and nutrient cycling processes in forest systems. By ensuring the retention of multiple species of CWD in various decay stages while forests develop and change, a wider spectrum of fungal biodiversity would be supported. We found nutrient concentrations in CWD (specifically Mn and K), total carbon, C/N ratio, carbon mineralization, and moisture content, were the best predictors of change in fungal communities within CWD. Further research to improve our understanding of these relationships is important, as fungal communities retain and, subsequently, release nutrients from CWD into the soil environment via decomposition, influencing ecosystem function. As noted above, retention of different species of CWD in various stages of decay in forest stands can also be important in forest carbon management, as we found that hardwood CWD had faster decomposition rates (microbial C mineralization) than conifer species, and late decay stage CWD across all species decomposed faster than CWD in early decay stages. This collective information will aid in developing strategies to determine the quality and quantity of CWD necessary to maintain carbon stocks in managed boreal forests.

## Research Data

Raw paired-end Illumina reads were submitted to the National Center for Biotechnology Information (NCBI) Short Read Archive (SRA) under BioProject ID PRJNA726337. Key infiles and scripts used to analyze the data and create figures are provided on GitHub at https://github.com/saskiahart.

## Acknowledgments

We would like to thank Kerrie Wainio-Keizer, Mathieu Levesque, and Alissa Ramsay for assistance in sample collection. Thank you to Susan Bowman, Kristine Pinkney, Derek Chartrand, and Daniel Doucet for assistance in laboratory analysis. Thank you to Emily Smenderovac for assistance in data analysis. Thank you to the Government of Canada through the Genomics and Research and Development Initiative (GRDI) Metagenomics-based Ecosystem Biomonitoring (EcoBiomics) project, the Ontario Ministry of Natural Resources and Forestry, and Laurentian University for financial support. The authors declare no conflict of interest.

## Supporting Information

**Table S1**: **Classification used for dividing CWD logs into different stages of decomposition in the field (Renvall, 1995; Sollins, 1981).**

**Table S2: Summary of bioinformatic processing.** *fITS7 - fITS4R ∼ 275 bp after removing primers, only 1 degenerate position,**fITS9 - fITS4R ∼ 298 bp after removing primers, with 2 degenerate positions.

**Figure S1. Rarefaction curves were used to assess sequencing depth. The horizontal line represents the 15^th^ percentile. Any further ESV based analysis used included at least 30,000 reads. Any samples with less than the 5^th^ percentile reads were removed from further analyses (3 samples removed).**

**Table S3. Pearson’s correlation coefficients (r values) associated with AICc model selection on the first two Principal Coordinate axes by wood physical and chemical parameters. Variables with a r >0.7 are highly colinear and are in bold.**

**Table S4. Analysis of variance (ANOVA) results with coarse woody debris characteristics as the dependent variables and stand, site (stand), tree species, and decay class as the independent variables (n=180). Significant values (p<0.05) are in bold.**

**Table S5. Analysis of variance (ANOVA) results with alpha diversity metrics as the dependent variables and stand, site (stand), tree species, and decay class as the independent variables (n=59). Significant values (p<0.05) are in bold.**

**Table S6. Results of a permutational multivariate analysis of variance (PERMANOVA) using binary Bray-Curtis dissimilarities. Comparisons were conducted within stand (strata= Stand Development Stage) and run with 999 permutations. Significant values (p<0.05) are in bold.**

**Table S7. Results of a permutational multivariate analysis of variance (PERMANOVA) using binary Bray-Curtis dissimilarities to determine whether Sites vary with Stand Development Stage. Comparisons were run with 999 permutations. Significant values (p<0.05) are in bold**.

## Notes

### Competing Interest Statement

The authors have declared no competing interest.

